# Optimal dormancy strategies in fluctuating environments given delays in phenotypic switching

**DOI:** 10.1101/2022.10.25.513531

**Authors:** Andreea Măgălie, Daniel A. Schwartz, Jay T. Lennon, Joshua S. Weitz

## Abstract

Organisms have evolved different mechanisms in response to periods of environmental stress, including dormancy – a reversible state of reduced metabolic activity. Transitions to and from dormancy can be random or induced by changes in environmental conditions. Prior theoretical work has shown that stochastic transitioning between active and dormant states at the individual level can maximize fitness at the population level. However, such theories of ‘bet-hedging’ strategies typically neglect certain physiological features of transitions to dormancy, including time lags to gain protective benefits. Here, we construct and analyze a dynamic model that couples stochastic changes in environmental state with the population dynamics of organisms that can initiate dormancy after an explicit time delay. Stochastic environments are simulated using a multi-state Markov chain through which the mean and variance of environmental residence time can be adjusted. In the absence of time lags (or in the limit of very short lags), we find that bet-hedging strategy transition probabilities scale inversely with the mean environmental residence times, consistent with prior theory. We also find that increasing delays in dormancy decreases optimal transitioning probabilities, an effect that can be influenced by the correlations of environmental noise. When environmental residence times - either good or bad - are uncorrelated, the maximum population level fitness is obtained given low levels of transitioning between active and dormant states. However when environmental residence times are correlated, optimal dormancy initiation and termination probabilities increase insofar as the mean environmental persistent time is longer than the delay to reach dormancy. We also find that bet hedging is no longer advantageous when delays to enter dormancy exceed the mean environmental residence times. Altogether, these results show how physiological limits to dormancy and environmental dynamics shape the evolutionary benefits and even viability of bet hedging strategies at population scales.

## I. INTRODUCTION

Dormancy is a reversible state of reduced metabolic activity that can protect an organism against unfavorable environmental conditions including nutrient deprivation, toxins, or temperature [1]. Dormancy is found across the tree of life and includes sporulation by bacteria [2], seed production by plants [3], and hibernation by mammals[4]. Despite profound differences in the underlying physiological mechanisms, these examples each share the common feature of dimorphism in growth, reproduction, and survival between the active and dormant state. These examples also differ in the ways in which dormancy is initiated or terminated.

Transitioning in and out of dormancy can be initiated randomly, i.e., independent of the environmental state, or as a result of an environmental sensing mechanism. Examples of stochastic entry and exit to and from dormancy span a variety of organisms such as persister cells, soil microbes, and plants [5–7]. Plants have dormant seeds which germinate randomly, presumably to avoid transitioning during only periods that have brief windows favorable for growth [8–10]. A variety of microorganisms employ a similar strategy in which cells exit dormancy at seemingly random times which allows the population as a whole to sample a spectra of environmental conditions [11–13]. Stochastic entry and/or exit from dormancy is presumed to balance the benefits of growth in good environments while avoiding death in bad environments.

Random transitioning between active and dormant states is often described as a bet-hedging strategy. A bet-hedging strategy refers to the investment in multiple phenotypes to reduce the potential loss suffered in future harsh environments, thereby increasing long-term or geometric mean fitness. Early work on bet-hedging theory was motivated by analysis of optimal betting strategies in (human) gambling [14]. As derived by Kelly [14], the optimal betting strategy is one that decreases the variance of possible outcomes by maximizing the logarithm of the expected growth in wealth. The so-called ‘Kelly’ optimal criterion was used to show that the optimal phenotypic switching rate should match the stochastic rate of environmental switching so as to to maximize the long-term Malthusian growth rate of replicating cells [[15]]. This approach has also been applied to compute the optimal life history outcomes for bacteriophage infections, given an analogy between dormancy and prophage integration and between active states and phage lysis of hosts [16].

Prior modeling efforts have tended to assume instantaneous switching between phenotypes when identifying the optimal rates of dormancy initiation and/or termination. In reality, transitioning between active and dormant can take a substantial amount of time, comparable to or even longer than an individual’s generation time. For example, spore-forming *Bacillus subtilis* has a generation time of 0.5 hours in rich media but it takes an individual bacterium several hours to initiate and transition into a dormant state [17, 18]. For other types of microbial dormancy, such as quiescence in persister cells or tumor cells, the time to reach dormancy is not as clearly defined, though it is often on the order of a replication cycle given the necessary changes in morphology and gene expression [19]. Hence, the time to reach dormancy varies by organism but still poses a significant opportunity cost. Depending on environmental fluctuations, delays in phenotypic switching could change the benefits and risks associated with a bet-hedging strategy.

In this paper, we extend the framework of bet-hedging models to explore the dynamic consequences of delays in reaching dormancy in both stochastic and predictable environments. Using Kelly’s insight of maximizing the expected logarithmic wealth we optimize for the Lyapunov exponent or equivalently for the expected long-term growth rate of the population. By using a multi-state Markov chain to simulate environments we can vary environmental simulations from memory-less to periodic-like. Consistent with previous work, we find that when phenotypic switching is instantaneous then optimal transitioning scales inversely with mean environmental residence time. However, when transitioning to dormancy takes additional steps and thus requires more time, the magnitude of optimal switching drops dramatically. Assessing bet hedging in the context of realistic delays and correlated environments can help deepen understanding of the drivers and constraints on the evolution of dormancy in biological systems.

## II. METHODS

### A. Summary

We propose a discrete stochastic model inspired by bet-hedging theory to compute the long-term fitness of a population in stochastic environments with two states: good or bad (see Fig. 1). Individuals can be active, dormant, or in transition from an active to a dormant state (Fig. 1 B). In this model, dormancy represents a metabolically inactive state invulnerable to environmental stressors, hence dormant individuals do not reproduce or die. Active individuals reproduce during good times, but die at a constant rate during bad times. Transition states represent the biological stages required to reach a fully developed dormant state. Therefore individuals in transition states are unable to reproduce but are also not protected from mortality and die during unfavorable environmental times. The transition states occur only from active to dormant states, with the inverse switch from dormant to active always taking one time step. This asymmetry is inspired by the relatively long period of generation of bacterial spores relative to their reactivation [18].

**FIG. 1:**
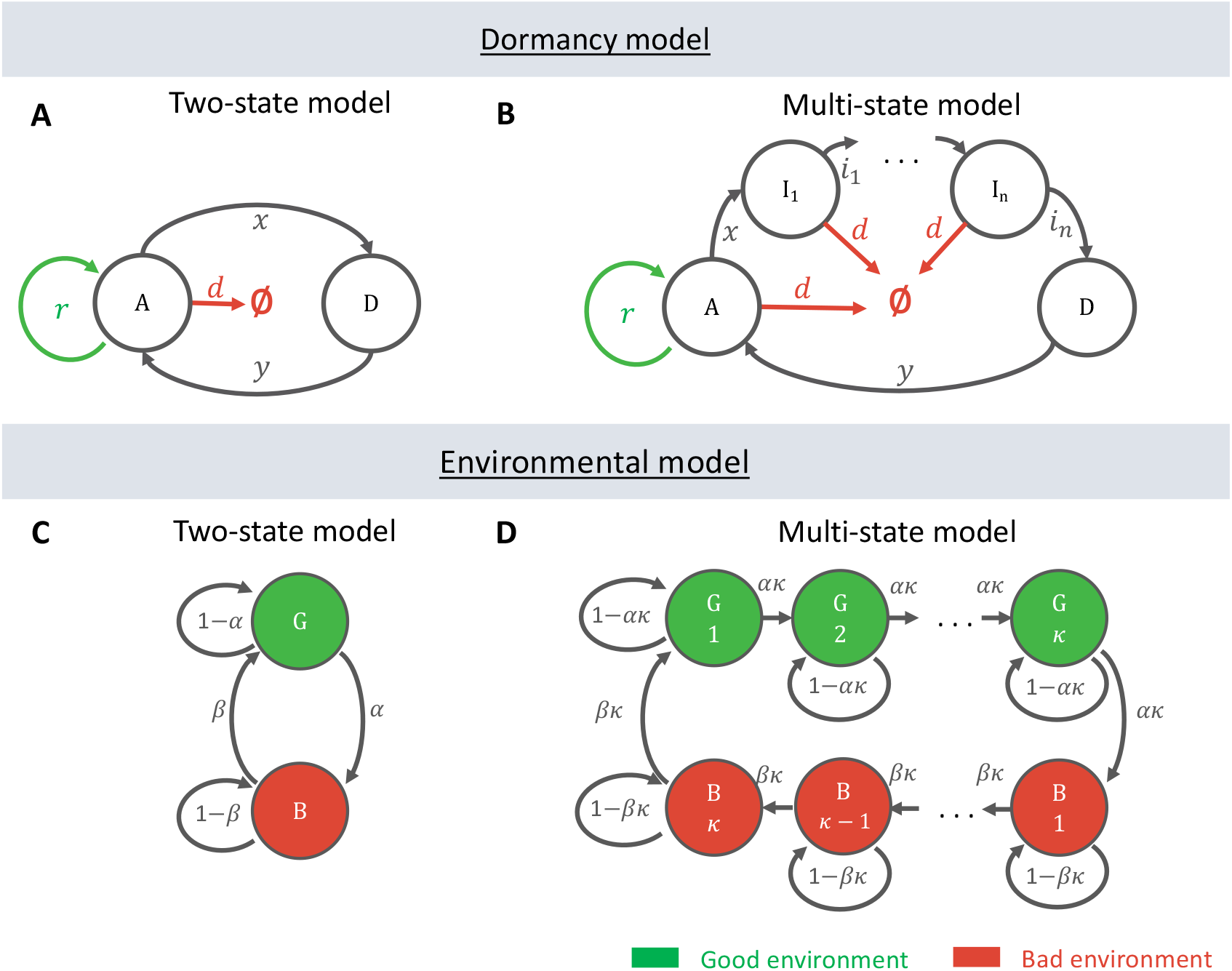
Schematic of dormancy and environmental models. **A and B** Compartmental model of active individuals A and dormant individuals D. Initiation probability is *x* and resuscitation probability is *y*. The green arrows represent growth during good times and red arrows death during bad times. Active individuals grow at rate *r* during good times and die at rate *d* during bad times. Dormant individuals are not affected by environmental conditions. **A.** Classical two-state dormancy model where transitioning between A and D is done in one time step. **B.** Extended multi-state model where transitioning between A and D involves passing through *n* transition states *I*_1_, *I*_2_, …, *I_n_* with probabilities *i*_1_, *i*_2_, …, *i_n_*. All transition states are vulnerable to bad environments, but cannot grow during good environments.**C.** Environments switch between good (G) and bad (B) based on a two-state Markov chain with switching probabilities *α* and *β*. **D.** Environments switch between good (G) and bad (B) by passing through all intermediate states *G*_1_, *G*_2_, …, *G_κ_* and *B*_1_, *B*_2_, …, *B_κ_*. There are *κ* good and *κ* bad states and the probability to advance through each state is *ακ* and *βκ*, respectively. This is a multi-state Markov chain and is an extension of the model in panel A.

We use a Markov chain to model the environmental conditions (Fig. 1 C, D) and to vary the correlation between time steps. We refer to the duration during which environments remain constant as the good or bad residence time. Because we choose the environmental parameters such that the residence time distribution is the same for good or bad environments, we use the term residence time to refer to both the good and bad times. By varying the switching rates and the number of intermediate states in the extended environment model we can control the mean and shape of the residence time distribution. The population and environmental models are coupled as the reproduction and death rates in the dormancy model are dependent on the environmental state.

### B. Dormancy model

We consider a two-state model in which individuals transition between active and dormant with initiation probability *x* and resuscitation probability *y* (Fig. 1 A, B). Active individuals grow at rate *r* during good times, but die at rate *d* during bad times. Dormant individuals are sheltered during bad times and cannot reproduce during good times. The transition states between active and dormant represent the delay it takes for active individuals to reach full refuge in dormancy (Fig. 1 B). Individuals in transition states do not reproduce but are susceptible to bad environments just like active individuals. These transition states are labeled *I*_1_, *I*_2_, …, *I_n_* with corresponding transitioning probabilities *i*_1_, *i*_2_, …, *i_n_*. Given a stochastic sequence of environmental conditions {*E*_1_, *E*_2_, …, *E_n_*} we can write a set of equations describing the number of active, transitioning, and dormant individuals as follows:

<u>Two-state model</u>

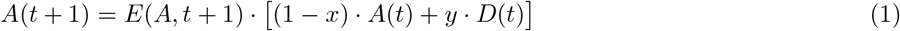

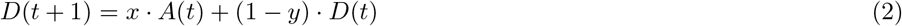

<u>Multi-state model</u>

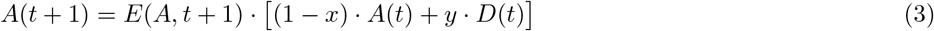

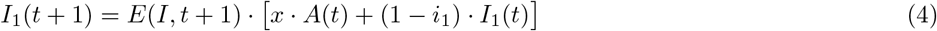

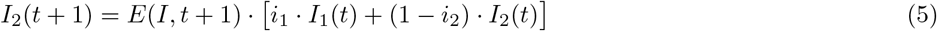

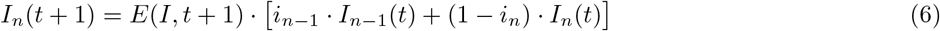

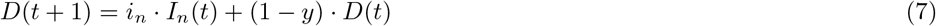

where *E*(*X, t*) gives the multiplication factor for state *X* ∈ {*A, I*} based on environmental condition *E_t_* as follows:

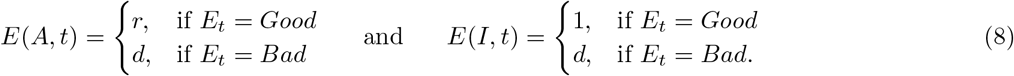

The conditions for *E*(*X, t*) are shown in Fig 1 A and B through the red and green arrows.

In this work we set *i*_1_ = *i*_2_ = … = *i_n_* = 1 meaning *n* transition states add a delay of n time steps to go from active to dormant. It is assumed that individuals in transition states cannot reproduce during good times, but are vulnerable to bad times and die at the same rate as active individuals. The values for the other parameters such as *r,d* and *n* are shown in table I.

**TABLE I:** Model variables

| Variable name | Description | Interpretation | Value |
| --- | --- | --- | --- |
| $\tau = \alpha^{-1} = \beta^{-1}$ | Mean residence time | Inverse environmental switching rate | [2, 4, 6, 8, 10, 20] |
| $\kappa$ | Minimum residence time | Environmental predictability | [1, 2, $\tau-2$, $\tau-1$] |
| $n$ | Number of dormancy transition states | Delay to reach dormancy | [0, 1, 2, 4, 6, 8, 10] |
| $x$ | Dormancy initiation probability | Bet-hedging investment | $0 \leq x \leq 1$ |
| $y$ | Dormancy resuscitation probability | Bet-hedging investment | $0 \leq y \leq 1$ |
| $d$ | Death rate | Bet-hedging risk | 0.5 |
| $r$ | Growth rate | Bet-hedging reward | 2 |

### C. Environmental model

Environments change between good and bad following a Markov process shown in Fig. 1, where the baseline model in Fig. 1 C represents a two-state Markov chain with switching between good and bad states and the extended model in Fig. 1 D is a multi-state Markov chain with additional states. These additional states are good or bad and individuals grow or die based on the same rules as in the two-state model. By introducing these intermediate states, different types of residence time distributions in good/bad states can be achieved. If we denote with *E_t_* the environment at time *t*, we can write the equations for the Markov chains in Fig. 1 as follows:

<u>Two-state model</u>

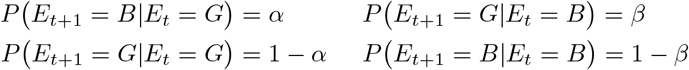

<u>Multi-state model</u>

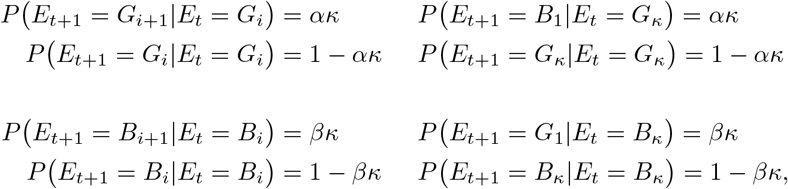

where *G* and *B* are the good and bad environmental states in the two state model and *G_i_* and *B_i_* are the *i^th^* good and bad state in the multi-state model. Both Markov chains have a steady-state distribution

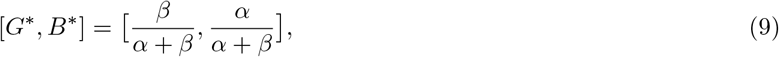

meaning that on average the environment will be good 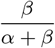 of the time and bad 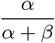. We maintain the fraction of bad times equal to the fraction of good times by setting 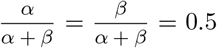, although the ratios can be changed independently between 0 and 1. We can vary *α* = *β* between 0 and 1 to obtain fast switching environments for higher values or slow switching environments for lower values.

In the two-state model, the distribution of environmental residence time is exponential with mean *α*^−1^ and *β*^−1^ respectively. For simplicity, we set the number of intermediate states to be the same in the good and bad states *κ*. Because *α* = *β*, let *τ* be the average length of time environments remain constant. As we increase the number of intermediate states *κ*, i.e. delays, in the multi-state model, the distribution of environmental residence time becomes more narrow but maintains the same mean *τ* (Fig. 2). Note that minimum residence time *κ* is anti correlated to the variance of residence time. When *κ* increases environments become more predictable and the distribution of residence time becomes more narrow (Fig. 2). We distinguish the following two parameters which we will use from now on to describe environmental conditions:

i. Mean residence time or the inverse of environmental switching rate *τ* = *α*^−1^ = *β*^−1^
ii. Minimum residence time or the number of intermediate environmental states *κ*.

**FIG. 2:**
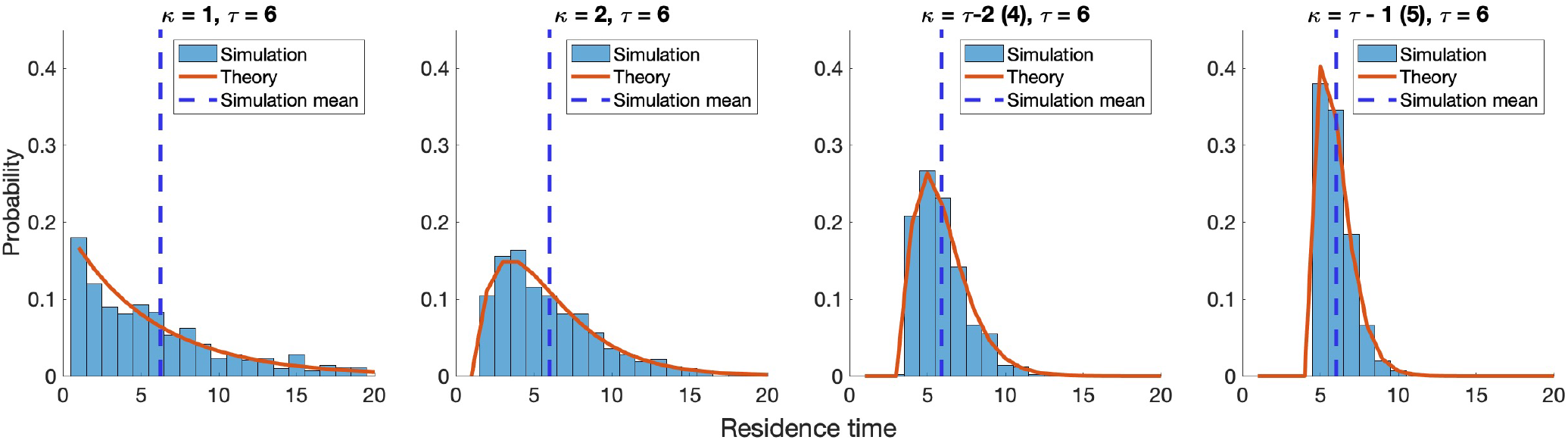
Probability density function of environmental residence time. Each of the four plots shows the PDF of residence time with the same mean residence time parameter *τ* = 6 and different values of minimum residence time *κ* between 1 and 5. Simulations of environments have 5000 time steps and are shown in the blue histogram. The mean residence time of the simulated environments is shown with a dotted blue line. A theoretical function computing the PDF based on the Markov process in Fig. 1 D is shown in orange.

The multi-state model includes a key constraint. Because all additional states need to be traversed to switch environments, we restrict our attention to models where *τ* > *κ*.

Given that *τ* = *α*^−1^ = *β*^−1^, *κ* has an upper limit of *α*^−1^ = *β*^−1^ and when that limit is reached environments are no longer stochastic. If the mean and minimum of the residence time distribution are equal, environments remain constant for exactly *τ* = *κ* time steps and are thus periodic. In this work we use *τ* ∈ {2, 4, 6, 8, 10, 20} and *κ* ∈ {1, 2, *τ* − 2, *τ* − 1} as described in table I. Additionally given *τ* > *κ*, not all pairs of (*τ, κ*) are feasible. The unfeasible cases, e.g. *τ* = 2 and *κ* = 2 are shown as ‘unfeasible’ in figures.

### D. Expected Lyapunov exponent

To compute the expected logarithmic growth rate, we use the Lyapunov exponent formula as a proxy for fitness:

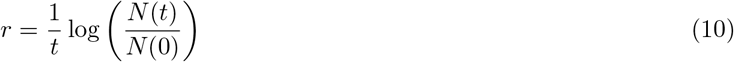

where *N* (*t*) = *A*(*t*) + *I*_1_(*t*) + … + *I_n_*(*t*) + *D*(*t*) in the multi-state model and *N* (*t*) = *A*(*t*) + *D*(*t*) in the two-state model. Generally *N* (*t*) represents the total size of the population at time *t*. A value of *t* = 500 time steps is used. We compute the Lyapunov exponent 1000 times for different simulations of the environment and take the mean to obtain the expected value.

### E. Code availability

All simulations were carried out in MATLAB v 2020a. Scripts are available on Github at https://github.com/WeitzGroup/Bet hedging dormancy.

## III. RESULTS

### A. Population dynamics and fitness change based on dormancy transitioning probabilities

We first evaluate how populations with different initiation and resuscitation probabilities grow and die in a stochastic environment using the two-state dormancy model. A stochastic environment of 50 time steps is simulated based on the Markov model in Fig. 1 A with values of *α* = *β* = 0.5. The population dynamics are shown in Fig. 3 A for three values of transitioning probabilities between 0 and 1. Populations grow during good times (green shading) and die during bad times (red shading). Each of the different populations has a fitness value calculated via the associated Lyapunov exponent shown in Fig. 3 B. Fig. 3 B also shows the Lyapunov exponent for a range of transitioning probabilities between 0 and 1 and note that the population with *x* = *y* = 0.5 yields the highest fitness.

**FIG. 3:**
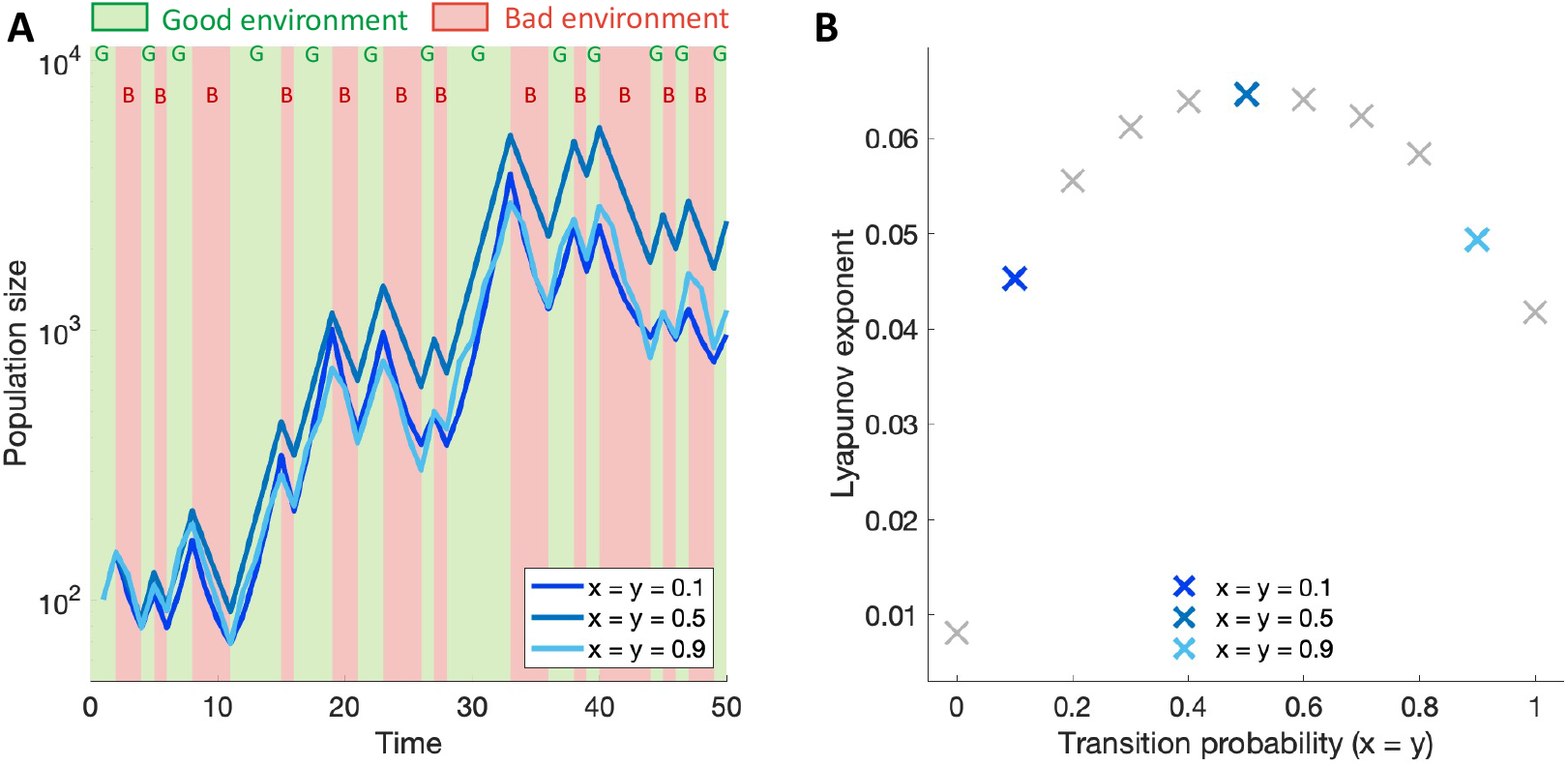
Population dynamics and Lyapunov exponent in the two-state dormancy model. **A.** Three populations with different initiation *x* and resuscitation *y* probabilities are shown in blue lines and environmental conditions are shown in red and green shaded areas which are also labeled G or B. The environmental and dormancy models used here are the two-state models from Fig. 1 panels A and C, respectively, with *τ* = *α*^−1^ = *β*^−1^ = 2 . **B.** The corresponding Lyapunov exponent for each of the blue lines in panel A is shown for *x* = *y* transitioning probabilities on the x-axis. The formula used to compute the Lyapunov exponent is shown in Eq. (10).

By varying (*x, y*) ∈ [0, 1]^2^ we generate a fitness map based on the initiation probability on the x-axis and resuscitation probability on the y-axis (Fig. 4 enlarged heatmap). This fitness map is concave and has a global maximum (*x*\*, *y*\*) = (0.5, 0.5) shown with a red mark. The pair (*x*\*, *y*\*) = (0.5, 0.5) are the optimal transitioning probabilities which maximize the fitness of the population given parameters *τ* = 2, *κ* = 1 and delay *n* = 0. The optimal solution can also be computed analytically by expanding the Lyapunov exponent and finding that the highest fitness is obtained when 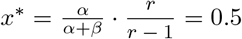 [16]. This analytical approach however does not hold for any other values of *n*, *κ* or *τ*, hence we need a computation approach to compute *x*\*, *y*\* for the remaining conditions. Fitness maps are shown in the rest of Fig. 4 for different values of mean residence time *τ* and minimum residence time *κ*. Notice that all the fitness maps in Fig. 4 are concave, hence we obtain an optimal pair of dormancy transitioning probabilities for each value of *τ* ∈ {2, 3, 4, 5, 6, 7} and *κ* ∈ {1, 2, 3, 4, 5, 6}. These results suggest that the fitness of a population is smooth with respect to the initiation and resuscitation probability and monotonically decreases as (*x, y*) pair is further away from the optimal value.

**FIG. 4:**
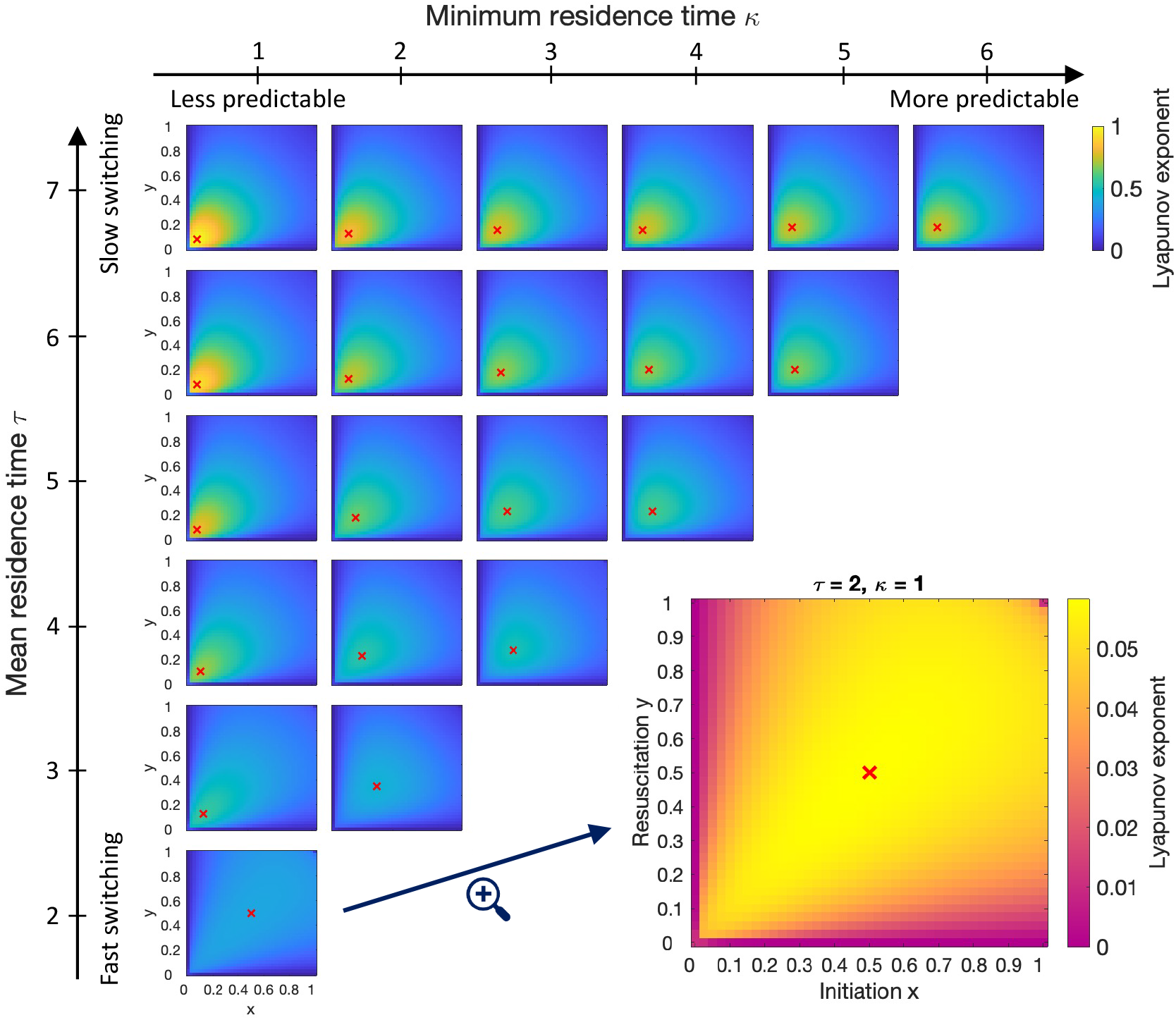
Fitness maps for no delays dormancy when varying mean and minimum residence time. Each heatmap shows the Lyapunov exponent based on the initiation probability on the x-axis and resuscitation probability on the y-axis. The maximum fitness is marked with a red x. The simulations are done for mean residence time *τ* ∈ {2, 3, 4, 5, 6, 7}, minimum residence time *κ* ∈ {1, 2, 3, 4, 5, 6} and no delays between active and dormant states (*n* = 0). Because *τ* ≥ *κ* the bottom right part of the figure is not feasible. Instead a magnified version of the *τ* = 2, *κ* = 1 heatmap is shown. The values for transitioning probabilities are between 0 and 1 with an increment of 0.025 and a total of 500 runs is used for each heatmap. In each case, the Lyapunov exponent shown is the average of the exponent for each run.

### B. Optimal transitioning in the two-state dormancy model based on residence time distributions

We evaluate the relationship between optimal transitioning probabilities identified in the prior section against the environmental residence time. Because transitioning into dormancy is instantaneous, we expect to find that optimal transitioning probabilities scale inversely with mean residence time [15]. We show the optimal pairs of dormancy transitioning probabilities identified in Fig. 4 as a function of *τ* in Fig. 5 A and as a function of *κ* in Fig. 5 B. First we note that the optimal initiation is equal to the optimal resuscitation, hence only one line is used to show *x*\* = *y*\*. Optimal transitioning decreases with *τ*, suggesting an inverse relationship between mean residence time *τ* and transitioning probabilities (Fig. 5 A). We also note that the optimal transitioning probabilities are closer to the theoretical inverse estimate as *κ* increases, i.e., this inverse relationship strengthens as environments become more predictable. For higher values of *κ* ∈ {*τ* − 2, *τ* − 1} the optimal transitioning approaches *τ* ^−1^. This observation is consistent with previous findings that optimal phenotypic switching rates are equal to the inverse of environmental times in the limit of sufficiently stable environments [15].

**FIG. 5:**
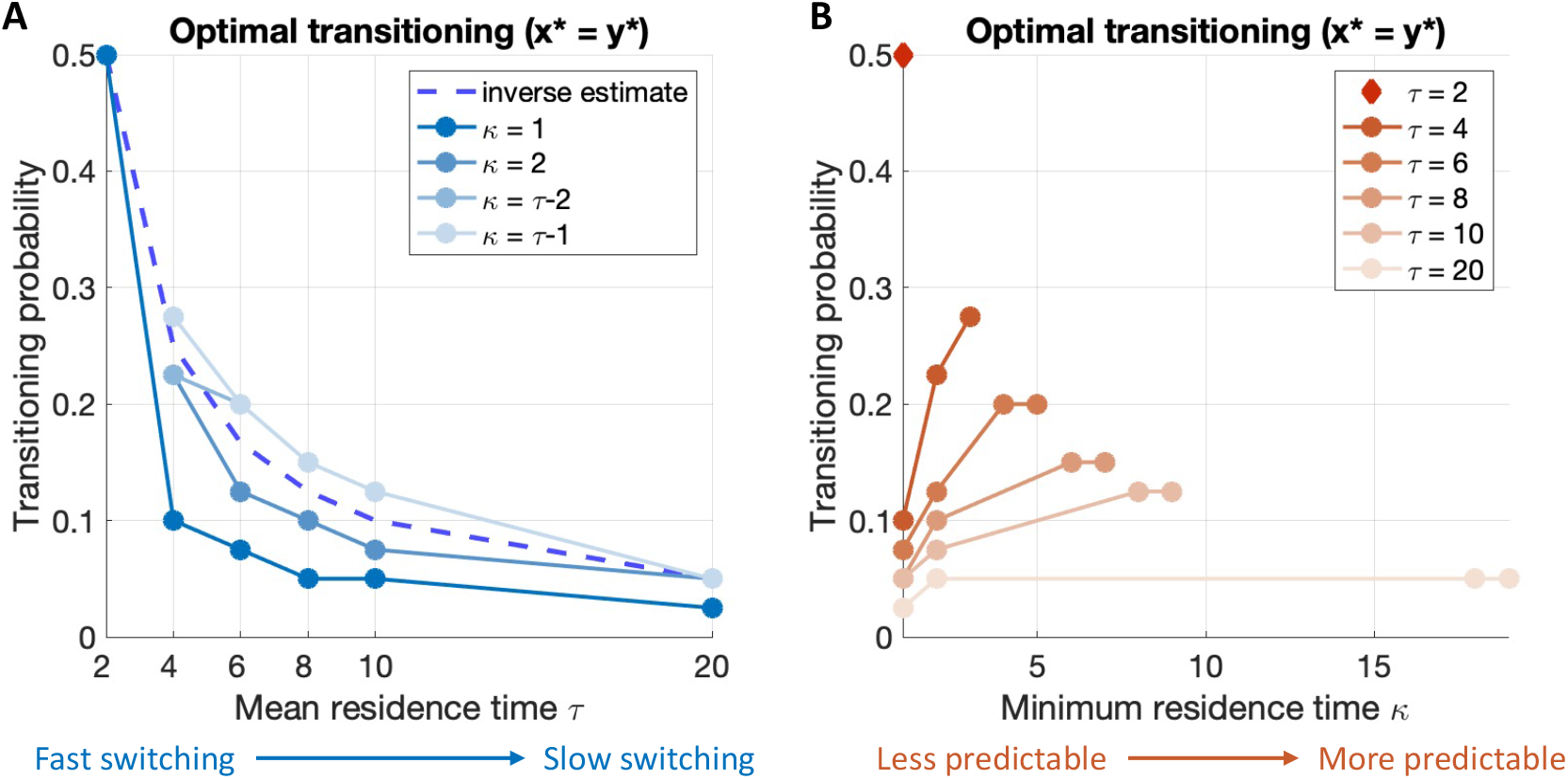
Impact of the mean residence time *τ* and minimum residence time *κ* on optimal transitioning with no delays. **A and B.** Optimal transitioning *x*\* and *y*\* are computed based on the location of the red x in Fig. 4 and shown against the mean residence time *τ* in panel **A** and against the minimum residence time *κ* in panel **B**. Note that *x*\* = *y*\*, hence one plot is representative of both optimal initiation and resuscitation probabilities. The optimal transitioning is shown for each value of *κ* in **A** and *τ* in **B**, with lower values of *κ* and *τ* being more opaque. The missing data points in both panels correspond to the case of *κ* ≥ *τ*, specifically when *τ* = 2 and *κ* ∈ {2, *τ* − 2, *τ* − 1}. The theoretical estimate from [15] is the inverse of the environmental times and is shown in a dashed line. The dormancy model considered is a two-state model with no delays when individuals transition from A and D.

We also evaluate the impact of the minimum residence time *κ* on the optimal transition probabilities (Fig. 5 B). As *κ* increases, the variance of environmental residence times decreases. Optimal transitioning probabilities increase as environments are more predictable and this observation is stronger for faster switching environments, i.e., for lower values of *τ*. Thus, in the absence of delays, the shape of the residence time distribution affects the optimal bet-hedging strategy only when environments switch frequently between good and bad times.

### C. Delays to reach dormancy reduce the fitness and optimal transitioning probabilities in a population

We next investigate how delays to reach dormancy affect a population by using the multi-state dormancy model introduced in Fig. 1 D. Whereas in previous sections we used a dormancy model with no delays, we compute the Lyanpunov exponents for delay *n* = 1 and show the results in Fig. 6. Note that Figures 4 and 6 are identical aside from variation in the delay time to dormancy *n*. When comparing the heatmaps with and without dormancy, we note that the addition of one time step to reach dormancy reduces the population fitness for all values of *κ* and *τ*. When comparing the enlarged heatmaps for *τ* = 2, *κ* = 1, the addition of one delay reduces the highest possible fitness by about 50% and the optimal transitioning probabilities decrease 10-fold from (*x*\*, *y*\*) = (0.5, 0.5) to (*x*\*, *y*\*) = (0.05, 0.05). Overall, the inclusion of delays to reach dormancy significantly reduces the long-term growth and leads to lower optimal transitioning probabilities throughout the parameter space explored in *τ* − *κ*.

**FIG. 6:**
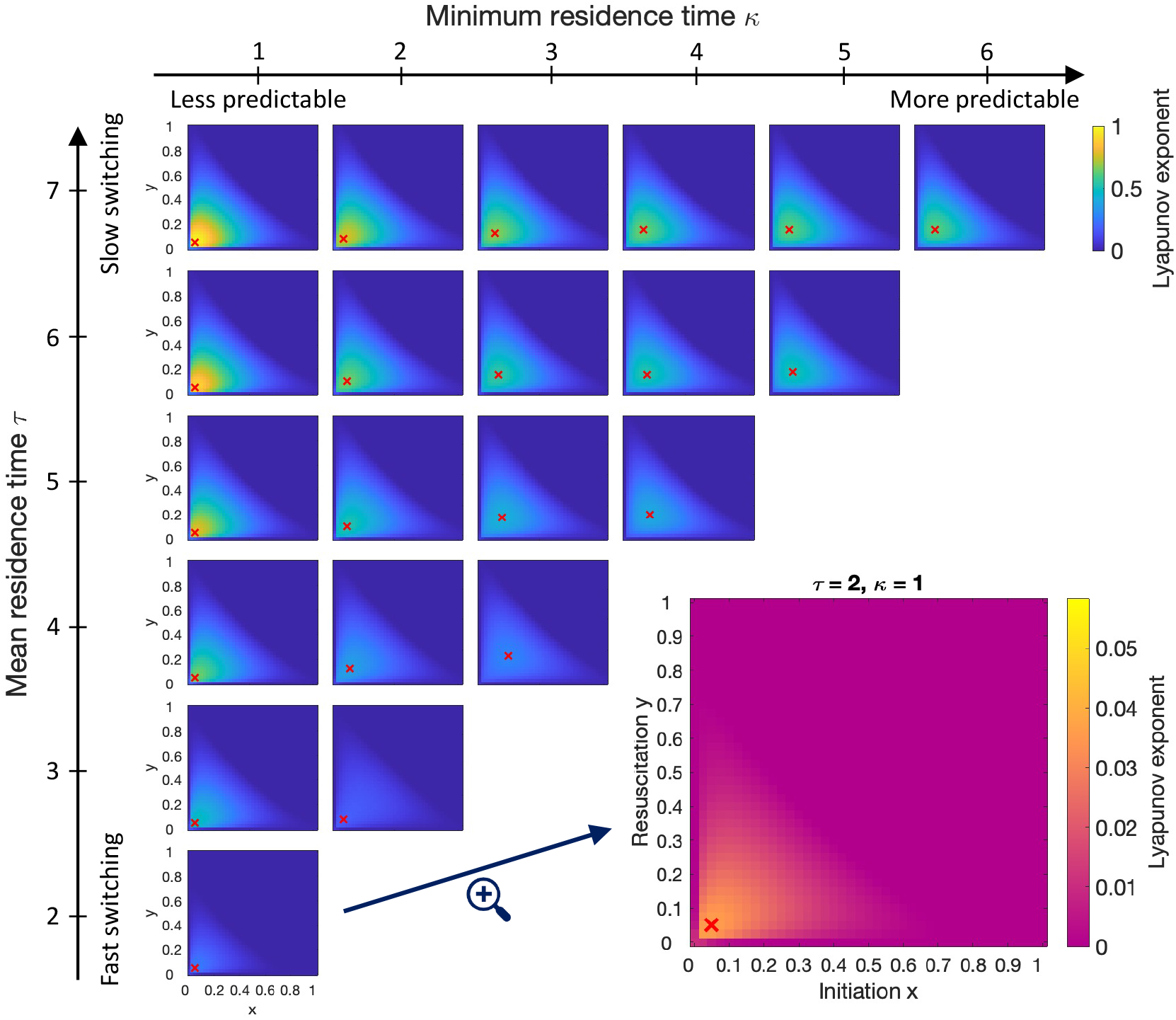
Fitness maps for multi-state dormancy model when varying mean and minimum residence time. Similar to Fig. 4, each heatmap shows the Lyapunov exponent based on the initiation probability on the x-axis and resuscitation probability on the y-axis. The only difference from 4 is that the dormancy model used here has one additional transition state (*n* = 1). The same values are used for *τ* and *κ* and a magnified version of the *τ* = 2, *κ* = 1 heatmap is shown in the unfeasible area. The values for transitioning probabilities are between 0 and 1 with an increment of 0.025 and a total of 1000 runs is used for each heatmap.

Building on this finding, we compute the optimal transitioning probabilities for other values of *τ, κ* and *n* and show that delays to reach dormancy *n* have a stronger effect on the fitness when the mean residence time *τ* is higher (Fig. 7). We note that the optimal initiation is equal to the optimal resuscitation within a small tolerance of 0.025 in the transitioning probability, likely due to stochastic effects, hence we show only one line for *x*\*. Both the initiation and resuscitation probabilities decrease as delays to reach dormancy increase until the optimal transitioning becomes *x*\* = *y*\* = 0 when the delays are too high (Fig. 7 A). This trend holds for all values of *τ*, however the reduction in optimal transitioning is more drastic for faster changing environments with lower mean residence time. The associated Lyapunov exponent also decreases as delays increase from almost 0.15 when *n* = 0 to ≈ 0.01 when *n* = 10 (Fig. 7 B). Altogether our findings suggest that increasing delays to reach dormancy decrease long-term fitness and decrease the optimal transitioning probability between active and dormant states - although the low levels of transitioning are retained for the optimal transitioning rates.

**FIG. 7:**
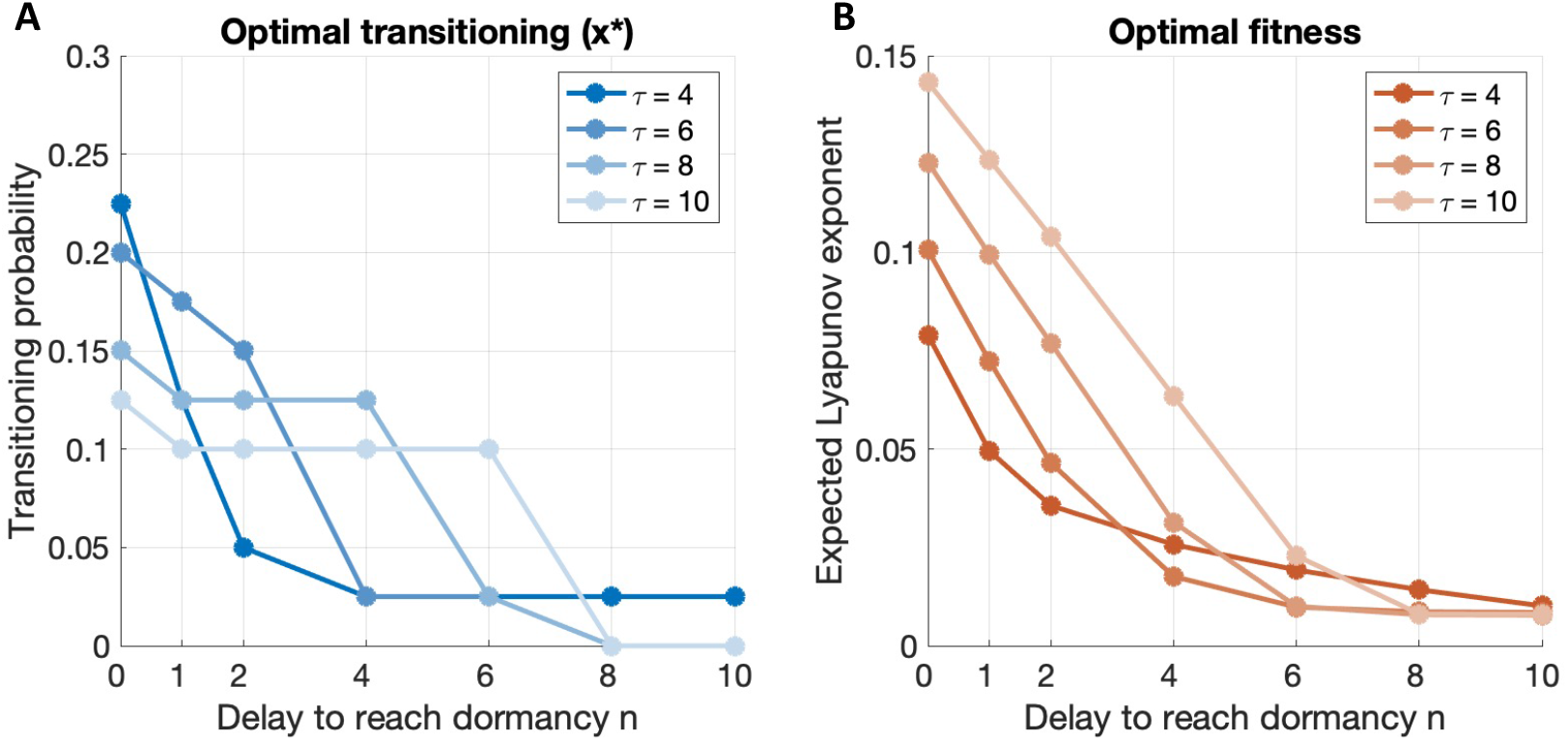
Effects of delays to reach dormancy on highest fitness and optimal transitioning probabilities. **A.** Optimal transitioning probabilities are computed for all values of *τ, κ* and delays and plotted against the delays to reach dormancy on the x-axis. Note that *x*\* = *y*\* within a small tolerance of 0.025, hence one plot is representative of both optimal initiation and resuscitation probabilities. The optimal transitioning is shown for several values of *τ*, with lower values of *τ* being more opaque. **B.** The corresponding fitness for the optimal transitioning probabilities is shown against the delay to reach dormancy on the x-axis. The fitness is shown for the same values of *τ* as in panel **A**, with lower values of *τ* being more opaque. Both panels use a value of *κ* = *τ* − 2 and the value of *τ* = 2 is omitted since it is unfeasible.

### D. Optimal transitioning increases with environmental predictability (*κ*) unless delays (*n*) are too large

In this last section we investigate the combined effect of residence time distribution and delays to reach dormancy on optimal transitioning and long-term fitness. To do so, we compute the optimal transitioning probabilities for all values of *τ, κ* and *n* in Fig. 8 and group the results in 4 panels based on minimum residence time *κ*. As the value of *κ* increases, environments become more predictable. As before, we find that *x*\* = *y*\* within a small tolerance of 0.025 for all values of *κ, τ* and *n*.

**FIG. 8:**
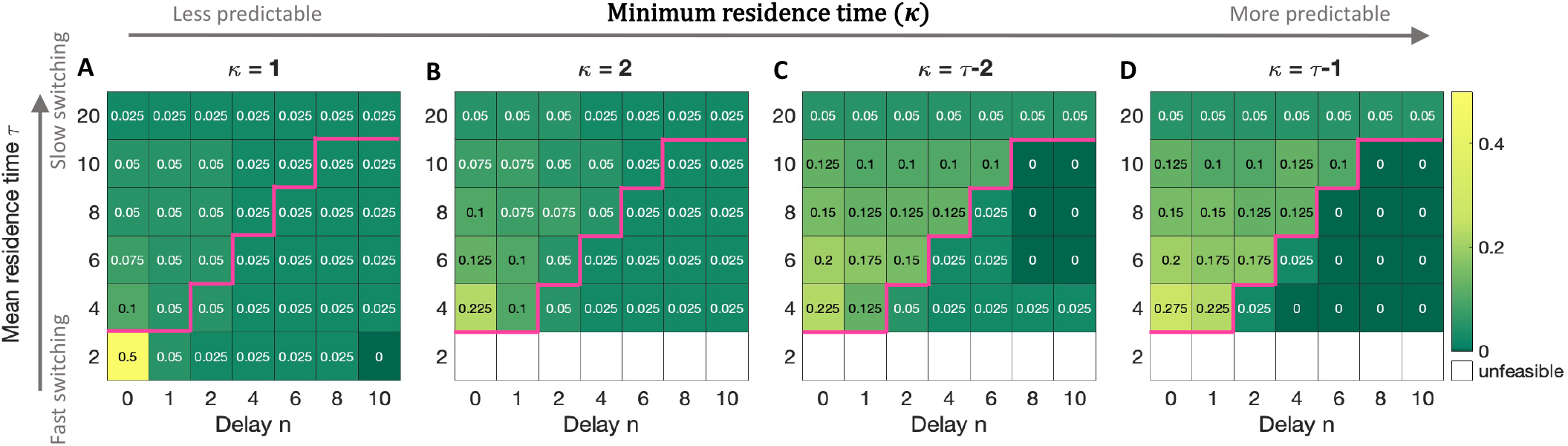
Optimal transitioning for all values of *τ, κ* and delays to reach dormancy. Each of the four maps shows the optimal transitioning probabilities based on the delay to reach dormancy *n* on the x-axis and mean environmental residence time *τ* on the y-axis. Each panel has a different environmental predictability with increasing values of *κ* from left to right. Given the constraint of *τ* > *κ*, unfeasible areas are left white. The magenta line in each panel represents the delimitation between *τ* + 2 and *n*: above the line *τ* + 2 > *n* and below *τ* + 2 ≤ *n*. The equation *τ* + 2 = *n* describes the situation in which the time to reach dormancy is most likely equal to the environmental residence time. The values for transitioning probabilities are between 0 and 1 with an increment of 0.025 and a total of 1000 runs is used to find each optimal value.

First we make note of results that are consistent with the ones from previous sections, but hold true for a larger range of parameters. Consistent with the results in section III C, optimal transitioning decreases as delays increase and this remains valid for all values of *τ* and *κ*. We also notice that, similar to the results discussed in section III B, transitioning probabilities are more dependent on the mean residence time *τ* as the value of *κ* is higher. Optimal transitioning probabilities tend to be equal to the smallest nonzero value for lower values of *κ* of 1 or 2. However when *κ* is higher there is a stronger correlation between *x*\* = *y*\* and both the environmental residence time and the delay to reach dormancy, *τ* and *n*, respectively. Indeed when *κ* = 1, all but four optimal transitioning values are equal to 0.05 or 0.025, whereas for *κ* = *τ* − 1 most intermediate values between 0 and 0.275 are obtained several times (compare Fig. 8 A vs. Fig. 8 D).

We further investigate how the optimal transitioning probabilities change with minimum residence time and observe a sharp switching behaviour based on mean residence time *τ* and delay to reach dormancy *n*:

1. Optimal transitioning probabilities increase with *κ* when *τ > n* + 2 (compare the top of each panel in Fig. 8).
2. Optimal transitioning probabilities decrease with *κ* when *τ* ≤ *n* + 2 until they reach 0 or 0.025 when *k* = *τ* − 1 (compare the bottom of each panel in Fig. 8).

We interpret these two observations as follows: as *κ* increases, *τ* = *n* + 2 represents the threshold when the environmental residence time is often equal to time to transition into dormancy. When *κ* = *τ* − 1, environments often remain unchanged for exactly *τ* −1 time steps. If *τ* = *n*+2, then environments remain constant for exactly *τ* −1 = *n*+1 time steps which coincides with the number of steps needed to transition between active and dormant (recall that delay 0 still takes one time step). If there is some uncertainty regarding how soon the environments will change, i.e., smaller values of *κ*, dormancy might still be beneficial even if the delays are longer than mean environmental residence times. However, when environments are highly correlated, the relationship between length of delays *n* and mean environmental residence times *κ* determines whether dormancy is a favorable strategy or not. In other words, if by the time dormancy is reached the environmental condition has changed, it would be more beneficial not to engage in dormancy at all.

## IV. DISCUSSION

Using a stochastic model of individual growth and dormancy coupled to fluctuating environments, we find that delays in transitioning into dormancy have a significant impact on the benefits of dormancy: both in reducing long-term population fitness and optimal transitioning probabilities. Delays in phenotypic switching decrease the benefits and increase the risk of investing in dormancy since transitory states are susceptible to harsh conditions. We find that the balance between the length of delays, mean residence time and minimum residence time determines how much and whether dormancy is a beneficial strategy. In particular, we find that dormancy is no longer beneficial when delays are consistently longer than environmental residence times, but dormancy can be maintained at a low level when environmental predictability is low. Thus while delays in phenotypic switching can have a drastic effect on optimal strategies environmental predictability alone can lead to the maintenance or loss of dormancy. These results show that all three components — environmental mean residence time, minimum residence time, and delay times — are essential in analyzing the benefits of dormancy as a bet-hedging strategy.

These findings build upon previous work on phenotypic switching which have prioritized the link between the mean environmental residence time and optimal transitioning probabilities. In classic bet-hedging theory, environmental conditions change randomly and are not correlated. In that case, corresponding to *κ* = 1, optimal transitioning probabilities match the theoretical prediction from [16]. When environments remain constant for longer periods of time and *κ* is closer to *τ*, we recapitulate the result that optimal transitioning probabilities should match the environmental switching probabilities [15]. By varying *κ* between 1 and *τ* we are able to see how the system behaves in between these limiting cases.

Our results highlight the importance of considering dormancy associated life-history traits when assessing optimal bet-hedging investment in a biological system. In the absence of a sensing mechanism, we predict that dormancy is favorable in unpredictable environments even when delays are relatively high. In contrast, in more predictable environments dormancy is beneficial only if environments remain constant longer than the time needed to transition into dormancy. For organisms with with long delays to reach dormancy, such as spore-forming bacteria or plants, our results suggest that stochastic entry and exit from dormancy is beneficial either in highly unpredictable environments or in environments that switch infrequently between states. For organisms with shorter delays to reach dormancy, such as those entering a quiescent state, randomly initiating dormancy may be beneficial regardless of environmental conditions, however higher optimal dormancy transitioning is obtained in more predictable environments.

We made several simplifying assumptions when modeling the link between dormancy, fluctuating environments, and long-term fitness. Environments are parameterized in a binary fashion as good or bad with fixed associated growth or death factors, whereas in reality environmental severity and organismal parameters can take on more than two values. Additionally, we assume that dormancy is purely stochastic without taking into account a responsive transitioning mechanism based on available resources or population density [20–23]. Indeed, our results suggest that when dormancy takes time to initiate, there may be a broad set of parameter combinations when responsive switching may outperform strictly bet hedging strategies [24]. Lastly, we note that this work does not explore a potential relationship between the delay to reach dormancy and the robustness of the dormant state against environmental stressors [25]. In reality, protective benefits of dormancy may vary with time of investment; hence it may be of value to explore potential trade-offs between different types of dormancy strategies when the benefits of dormancy also vary with the time to reach it. Using the framework presented in this paper could provide further insight into the trade-off between the quantity and quality of dormant individuals.

In summary, our findings extend previous efforts to analyze bet-hedging strategies by showing the importance of incorporating delays to reach dormancy. As we have shown, small changes in the mechanistic representation of dormancy and/or environmental conditions can have large effects on the benefits of particular bet-hedging strategies. Future work incorporating mixed strategies, direct competition between populations, and sensing-based initiation and exiting of dormancy could provide insights regarding the benefits and robustness of evolutionary strategies in fluctuating environments.

## Declaration of Competing Interest

The authors declare that they have no known competing financial interests or personal relationships that could have appeared to influence the work reported in this paper.

## Acknowledgments

We would like to thank Guanlin Li, David Demory, and Jacopo Marchi for feedback with the bet-hedging framework. This work was enabled by support from the National Science Foundation (DEB - 1934554 to JTL and DS, DBI - 2022049 to JTL and DEB - 1934586 to JSW and AM), US Army Research Office Grant (W911NF-14-307 1-0411 and W911NF-22-1-0014 JTL) and the National Aeronautics and Space Administration (80NSSC20K0618 JTL). JSW was supported, in part, by the Chaires Blaise Pascal program of the Ile-de-France region.

## Notes

### Competing Interest Statement

The authors have declared no competing interest.

